# Hierarchical annotation of eQTLs enables identification of genes with cell-type divergent regulation

**DOI:** 10.1101/2023.11.16.567459

**Authors:** Pawel F. Przytycki, Katherine S. Pollard

## Abstract

While context-type-specific regulation of genes is largely determined by cis-regulatory regions, attempts to identify cell-type specific eQTLs are complicated by the nested nature of cell types. We present a network-based model for hierarchical annotation of bulk-derived eQTLs to levels of a cell type tree using single cell chromatin accessibility data and no clustering of cells into discrete cell types. Using our model, we annotated bulk-derived eQTLs from the developing brain with high specificity to levels of a cell-type hierarchy. The increased annotation power provided by the hierarchical model allowed for sensitive detection of genes with multiple distinct non-coding elements regulating their expression in different cell types, which we validated in single-cell multiome data and reporter assays. Overall, we find that incorporating the hierarchical organization of cell types provides a powerful way to account for the relationships between cell types in complex tissues.

## Introduction

Context-specific regulation of gene expression is largely determined by noncoding cis-regulatory regions.^1,2^ These sequences encode information about the time, place, and quantity in which a gene will be transcribed allowing for tissue and cell-type specific regulation.^1,2^ While it is well established that genes are pleiotropic,^1,2^ the way in which regulatory elements specify the contexts in which a gene will be expressed is complex and not well understood.^2^

Expression quantitative trait loci (eQTLs) measure the association between genetic variants and gene expression across large cohorts. In order to capture changes in expression in specific tissues, large-scale efforts such as GTEx take expression measurements in tissues across many individuals.^3^ However, because many regulatory effects are cell-type specific, recent work has begun to identify cell-type specific eQTLs by taking massive amounts of single-cell gene expression measurements (scRNA-seq) across cohorts.^4–6^ Unfortunately, such approaches are not broadly feasible due to the high cost of single-cell sequencing.^6^ An alternative approach is to deconvolve bulk derived eQTLs into cell-type specific signatures based on scRNA-seq data for those cell types^7^ or based on single-cell chromatin accessibility of the region containing the genetic variant.^8,9^

Attempts to identify cell-type specific eQTLs are complicated by the nested relationships between cell types. While most analyses assume discrete cell types,^10^ complex tissues such as the brain contain many rare and highly correlated cell types and subtypes that are not entirely distinct from each other.^11,12^ Cell states further complicate discretization. Therefore, the typical approach used with single-cell sequencing data of first generating clusters of similar cells and then assigning cell type labels to whole clusters often fails to capture the true diversity of cell types.^11^ This in turn constrains the identification of the context in which genes are expressed from single-cell sequencing data to the most common and most distinct cell types.

Towards addressing these challenges, we present a network-based model for hierarchical annotation of bulk-derived eQTLs using single cell chromatin accessibility data (scATAC-seq). Our model explicitly takes into consideration the tree-based organizational principle underlying cell diversity,^13^ rather than treating cell type as a categorical variable, and scores bulk eQTLs at all levels of a cell hierarchy to best identify significant cell-type and subtype specific annotations. These scores are based on chromatin accessibility of the genetic variant for each eQTL across cell types. Because our model annotates the genetic variant rather than the associated gene, we allow for gene pleiotropy and associate each variant in a locus with the expression of the target gene in potentially unique cell types.

We applied our model to eQTLs from the developing human brain,^9^ a complex organ with many rare or nested cell types and subtypes.^14,15^ We annotated 5,889 of these bulk-derived eQTLs with high specificity to levels of a cell-type hierarchy. Based on these hierarchical cell-type labels, we identified 613 genes with multiple eQTL variants that are specific to distinct cell types that we term cell-type divergent eQTLs. Using multiome single cell accessibility and expression data,^16^ we confirmed that cells in which the given eQTL variant site is accessible express the linked gene in the multiple predicted contexts. Finally, we dissected the regulation of *FABP7* and *ICA1L*, two genes expressed in the developing brain with multiple cell-type divergent eQTLs, using Massively Parallel Reporter Assay (MPRA) data.^17^ We we observed that both genes have eQTL variants with independent regulatory effects in the developing brain, confirming that they are functional variants. Overall, our hierarchical method generated an annotation of bulk eQTL data that allowed for the discovery of divergent cell type regulation in an organ with a complex mixture of cell types.

## Results

### Hierarchical model for nested cell types

We developed a network-based hierarchical model to identify cell-type specific eQTLs in complex tissues with closely related and nested cell types (Figure 1a). Our model extends the existing CellWalkR model^18^ to take a cell-type hierarchy as input in addition to cell-type labels and scATAC-seq data. Briefly, the cell type hierarchy is taken as prior knowledge, and it is implemented as edges between leaf nodes that represent specific cell types and internal nodes that represent broader cell types higher in the hierarchy. The cell type nodes are then connected to nodes representing cells based on how well marker genes correspond to each cell’s chromatin accessibility, and cells are connected to each other based on the similarity of their genome-wide chromatin accessibility. A random walk with random restarts model of network diffusion is then run on this network to calculate how much information flows from each node to each other node. In particular, this includes the probability that a walk starting at each cell node ends at each cell type node as well as each internal node representing portions of the cell-type hierarchy.

**Figure 1.**
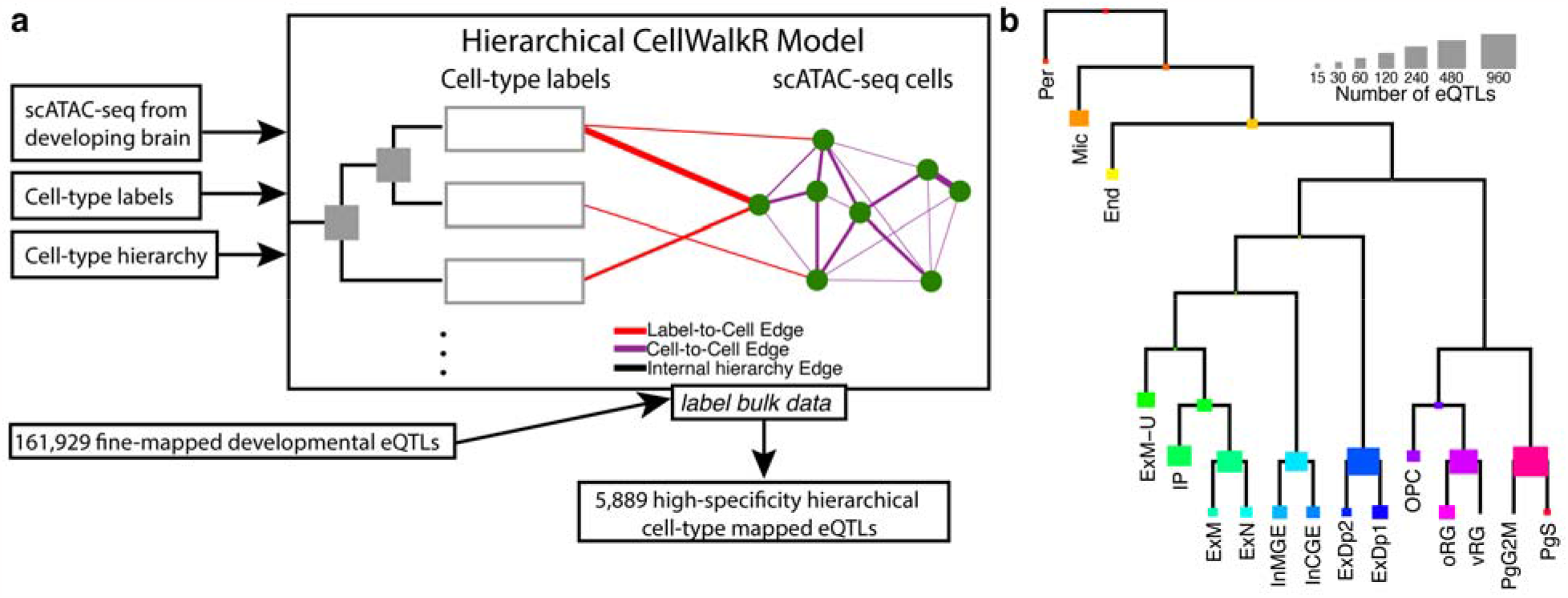
Hierarchical cell type mapping. **a**. The hierarchical model extends CellWalkR to take a cell-type hierarchy as an input in addition to scATAC-seq data and cell-type labels. The hierarchy is used to create internal nodes in the model that correspond to cell types higher in the hierarchy. This hierarchical model was used to label a large set of fine-mapped developmental brain eQTLs with high specificity. **b**. Count of how many developmental brain eQTLs are mapped to each hierarchical cell type.

Next, given a set of bulk-derived eQTLs, each eQTL is mapped to hierarchical cell types using the calculated flow of information and the chromatin accessibility of individual cells. For each eQTL, the location of the variant is intersected with the accessibility of each cell, and then the normalized cumulative flow from those cells is used to score each hierarchical cell type. This results in a score for each hierarchical cell type for each eQTL variant. In this way, eQTLs from bulk data can be mapped to a cell type tree. The cell type with the highest score can be used as a discrete labeling of the eQTL, or the scores across all cell types can be treated as a fuzzy (i.e., probabilistic) labeling.

### Annotation of eQTLs in the developing brain

We next applied this hierarchical model to label bulk eQTLs from the developing brain^9^ using scATAC-seq data from the mid-genstation telencephalon^19^ combined with a transcriptomics-based cell-type hierarchy derived from similar samples.^15^ Our model was able to label 5,889 eQTLs to hierarchical cell-types with high-specificity (z-score greater than two, see methods for details). These eQTLs mapped to a large variety of hierarchical cell types (Figure 1b), including both specific cell types (e.g. outer vs ventral radial glia) as well as higher level annotations (e.g. broadly neuronal).

For comparison, we annotated eQTL variants with a non-hierarchical version of the same model. We found that without hierarchical cell types, while the model was still able to label highly distinct cell types such as endothelial cells and microglia, it was unable to label similar or nested cell types such as different radial glia (Figure 2a, Supplementary Figure 1a and b). Only 1,252 eQTLs could be mapped to non-hierarchical cell types with high specificity, indicating a five-fold loss in annotation compared to using the cell type tree (Figure 2b). A less stringent threshold for specificity (z-score greater than one) annotates more eQTLs, but maps multiple non-hierarchical cell types to each eQTL (Supplementary Figure 1c), likely due to related cell types having highly correlated scores (Supplementary Figure 2). This indicates that a major advantage of using a cell type tree is its ability to account for highly correlated cell types.

**Figure 2:**
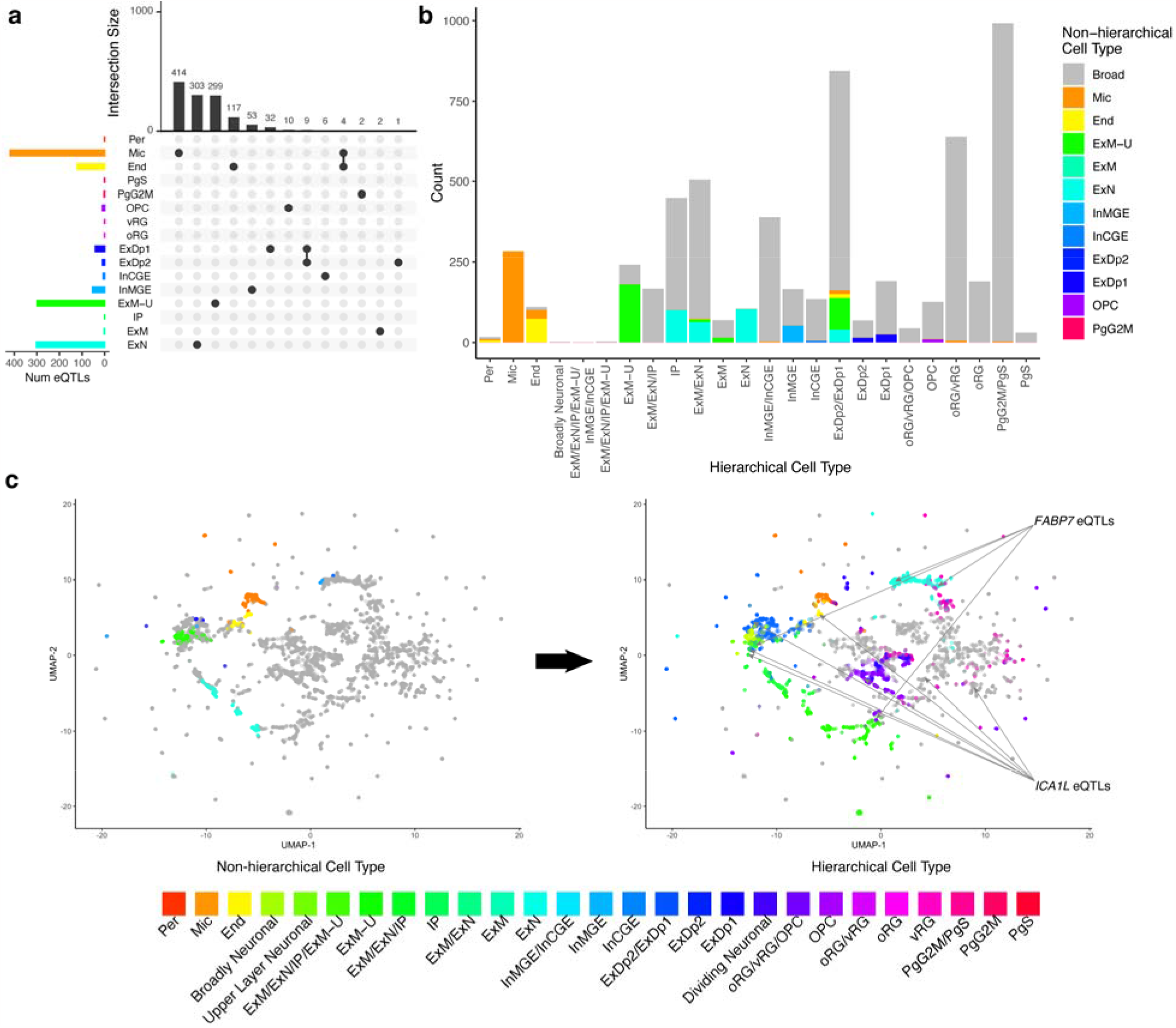
Hierarchical cell types provide improved labeling. **a**. Non-hierarchical high-specificity cell-type mapped eQTLs generally map to a single cell type, but they are biased towards very distinct or common cell types. **b**. Many eQTLs that could not be mapped to a specific cell type in the non-hierarchical model (“Broad,” shown in gray) receive hierarchical cell type labels (shown on x-axis). **c**. UMAP embedding of eQTLs labeled by non-hierarchical cell type (left) and hierarchical cell type (right) shows that a diverse set of previously unlabeled eQTLs can now be labeled. Due to this increased label diversity we can observe that some eQTLs linked to the same genes (e.g. *FABP7* and *ICA1L*) map to vastly different cell types.

As an orthogonal comparison, we overlapped eQTL variant sites with annotated broad cell-type specific peaks and enhancers^19^ and compared these annotations to our hierarchical and non-hierarchical eQTL cell type labels. Overall, we observed that our labels are consistent with these two sources of regulatory element annotation (Supplementary Figure 3). For example, we found that 93% of our hierarchically labeled eQTLs overlap cell type specific peaks. Non-hierarchical eQTL labels were also generally consistent with the annotations, but fewer of them overlapped cell type specific peaks and enhancers compared to hierarchical labeling. Together, these analyses validate our cell type labels and underscore the extra sensitivity provided by the cell type tree.

Using the label scores we calculated for each hierarchical cell type for each eQTL, we embedded the eQTL variants into two-dimensional UMAP space (Figure 2c). Consistent with the previous results, we observed a large increase in the coverage and diversity of hierarchical annotations of eQTLs as compared to non-hierarchical annotations. Hierarchically related cell types are located near each other, reflecting their relationships being modeled in the eQTL labeling process. Further, eQTL variants tend to cluster by cell type, rather than by the gene each variant is linked to. For example, the four eQTLs for the gene *FABP7* link to variants annotated to three different hierarchical cell types, and the six eQTLs for *ICA1L* are annotated to four different hierarchical cell types. These multi-cell-type annotations were not detected with the non-hierarchical model or by overlapping with cell-type specific peaks, emphasizing the need for hierarchical cell type annotation.

### Identification of cell-type divergent eQTLs

Given the hierarchical model’s increased ability to assign multiple distinct cell types to different eQTL variants linked to the same gene, we sought to identify all such genes. For each gene, we considered it to have cell-type divergent eQTLs if at least two eQTL variants linked to that gene were not the same cell type nor were they ancestors of each other in the original cell-type hierarchy. We also required that the full label scores for the eQTL variants were not similar to each other (see methods for details). We identified 613 genes with cell-type divergent eQTLs, the majority of which were linked to two distinct cell types though a few had three or more cell type annotations (Figure 3a). For comparison, only 88 genes with cell-type divergent eQTLs could be identified using the non-hierarchical model, only 320 using any overlaps with annotated cell-type specific peaks, and only 33 using overlaps with cell-type specific enhancers (Supplementary Figure 4). Thus, the higher sensitivity of our hierarchical model revealed a greater frequency of genes with eQTL variants that function in distinct cell types.

**Figure 3.**
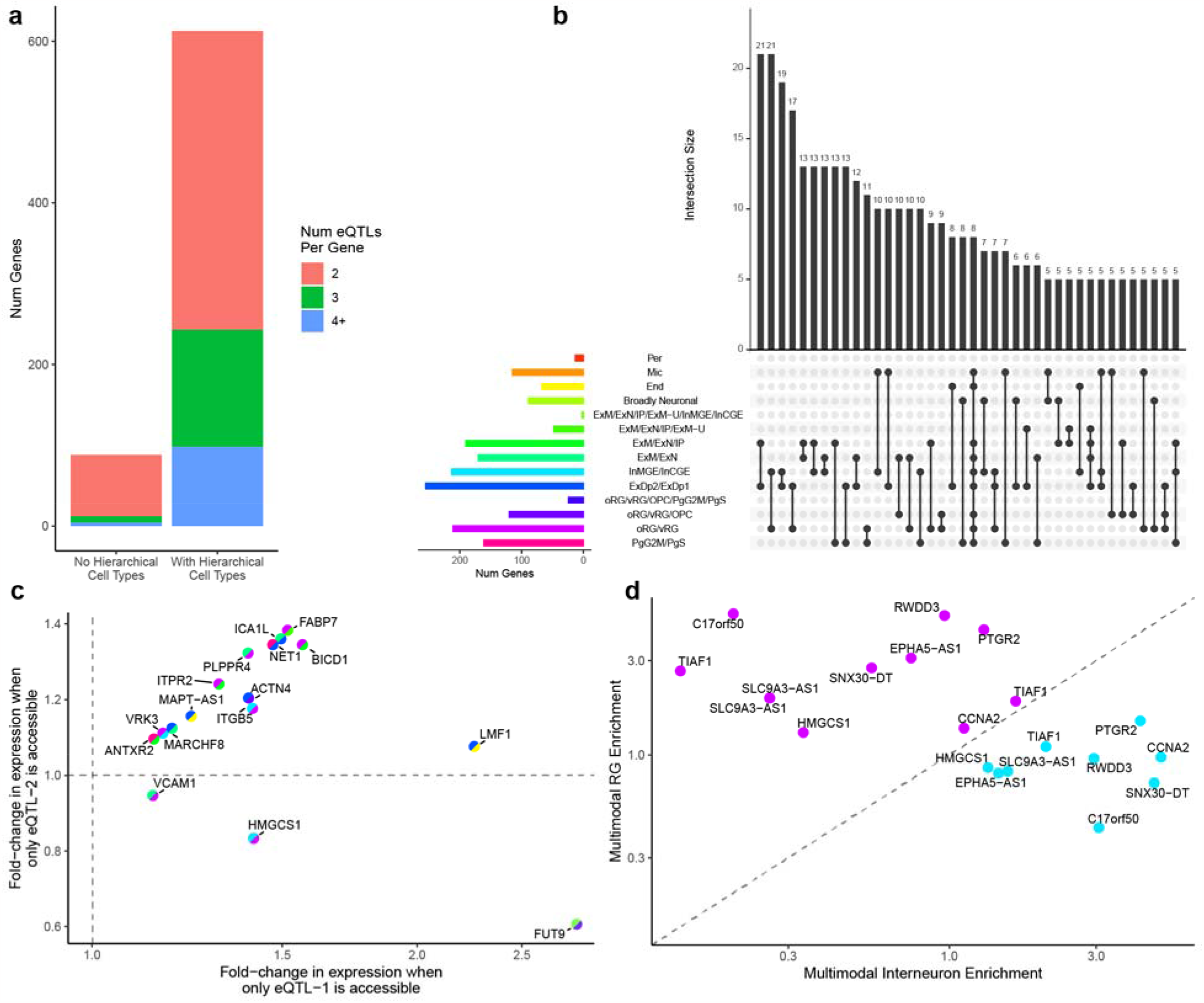
Cell-type divergent eQTLs. **a**. While only 66 genes have at least two eQTLs with distinct non-hierarchical cell types, the larger number of annotations we can make with the hierarchical model results in 613 genes having at least two distinct hierarchical cell types. **b**. An upset plot showing the most common divergent hierarchical cell types for eQTLs across genes. **c**. For highly expressed genes with cell-type divergent eQTLs, the gene is differentially expressed in cells where the first eQTL variant is accessible (x-axis) as well as in cells where the second eQTL variant is accessible (y-axis), as observed in jointly profiled multiome scRNA/ATAC data. The split colors of each point indicate the divergent hierarchical cell types following the key from panel b. **d**. For highly expressed genes with divergent Radial Glia (oRG/vRG, shown in magenta) and Interneuron (InMGE/InCGE, shown in teal) eQTLs, cells in which the respective eQTL variant is accessible are enriched for the matching cell type in the labeled scRNA-seq portion of jointly profiled multiome scRNA/ATAC data.

Taking a closer look at genes with cell-type divergent eQTLs, we find genes with eQTL variants corresponding to diverse combinations of cell types such as deep layer plus maturing excitatory neurons (21 genes) and interneurons plus radial glia (21 genes) (Figure 3b). For comparison, the non-hierarchical model almost exclusively identified cell-type divergent genes with eQTLs involving microglia and endothelial cells, two particularly distinct cell types, and generally failed to identify divergent eQTLs annotated to different types of neurons (Supplementary Figure 5). Additionally, we observed that the more cell-type specific eQTLs a gene has, the higher its expression entropy across cell types in scRNA-seq^16^ (Supplementary Figure 6). This indicates that the more different cell-type specific eQTLs a gene has, the more diverse the expression of that gene across cell types, consistent with these eQTLs providing cell-type specific regulation and making the gene more pleiotropic.

### Genes with cell-type divergent eQTLs exhibit cell-type specific regulation

In order to determine if cell-type divergent eQTLs directly contribute to cell-type specific expression, we looked at multiome measurements of scRNA-seq and scATAC-seq in the same cells in the developing brain.^16^ Given the sparse nature of multiome data, only 356 genes with cell-type divergent eQTLs could be tested in the multiome data. For each of these genes and each of their eQTL variants, we tested whether cells in which the variant was uniquely accessible (i.e. no other eQTL variant was accessible for that gene) the gene was differentially expressed relative to cells in which the eQTL was not accessible. We observed significant differential expression across multiple eQTLs for 30 of these genes (false discovery rate < 0.05, supplementary Figures 7 and 8). 16 of these differentially expressed genes were highly expressed (Figure 3c). Since accessibility and expression were determined using very sparse data per cell, we posit that the lack of significant differential expression for most eQTLs is influenced by low power. Overall we observe that changes in accessibility in cell-type divergent eQTLs lead to changes in expression in those same cells.

Next, we considered the cell type annotations of the scRNA-seq portion of the multiome data in order to determine if predicted cell-type divergent eQTL variants had cell-type specific accessibility. We observe that generally the scRNA-seq-based cell type annotations are enriched in the predicted cell type when the corresponding eQTL variant is accessible (Supplementary Figure 9). Furthermore, for highly expressed genes with cell-type divergent eQTLs, we detected an enrichment for corresponding cell types when each eQTL variant was uniquely accessible (Supplementary Figure 10). For example, looking at genes that have divergent eQTLs for radial glia and interneurons, we see that the eQTL variant is always more enriched in the predicted cell type when the corresponding variant is accessible (Figure 3d).

Finally, we looked for mechanisms of action for the change in expression. We found that of the 613 genes with cell-type divergent eQTLs, 222 had eQTL variants that disrupted binding sites of at least two different transcription factors (TFs) that are expressed in the corresponding cell type. Among these TFs, some are very specific to a single hierarchical cell type (e.g. *FOSB* and *JUND* for cycling PgG2M/PgS progenitor cells, *FOXP2* for newborn and maturing ExM/ExN/IP excitatory neurons) while some are disrupted in many cell types (Supplementary Figure 11). Furthermore, some TFs frequently co-occur as disrupted by cell-type divergent eQTLs (Supplementary Figure 12). For example, 13 genes have both an eQTL predicted to disrupt *SMAD2* binding in newborn and maturing excitatory neurons (ExM/ExN/IP) and an eQTL predicted to disrupt *JUND* binding in cycling progenitor cells. This supports the idea that one mechanism of gene pleiotropy is cell-type specific transcription factor binding.

### Cell-type divergent regulation of *FABP7* and *ICA1L*

The brain-related genes *FABP7* and *ICA1L* are both expressed in multiple cell types in the developing brain (Figure 4a). *FABP7*, which plays a role in the establishment of radial glial fiber,^20^ has four eQTLs that we mapped to hierarchical cell-types. Of the three variants that overlapped peaks in multiome scRNA/ATAC data, each was enriched for the corresponding scRNA-seq cell types annotation when the variant was accessible (Figure 4b, top). Furthermore, two eQTLs overlapped known enhancers, two overlapped predicted cell-type specific regulatory elements, and three disrupted different TFs that were expressed in the corresponding cell types, indicating a possible mechanism of action (Supplementary Figure 13). Taken together, this suggests that the expression of *FABP7* in different cell types may be driven by cis regulatory elements overlapping eQTL variants as annotated by our hierarchical model.

**Figure 4.**
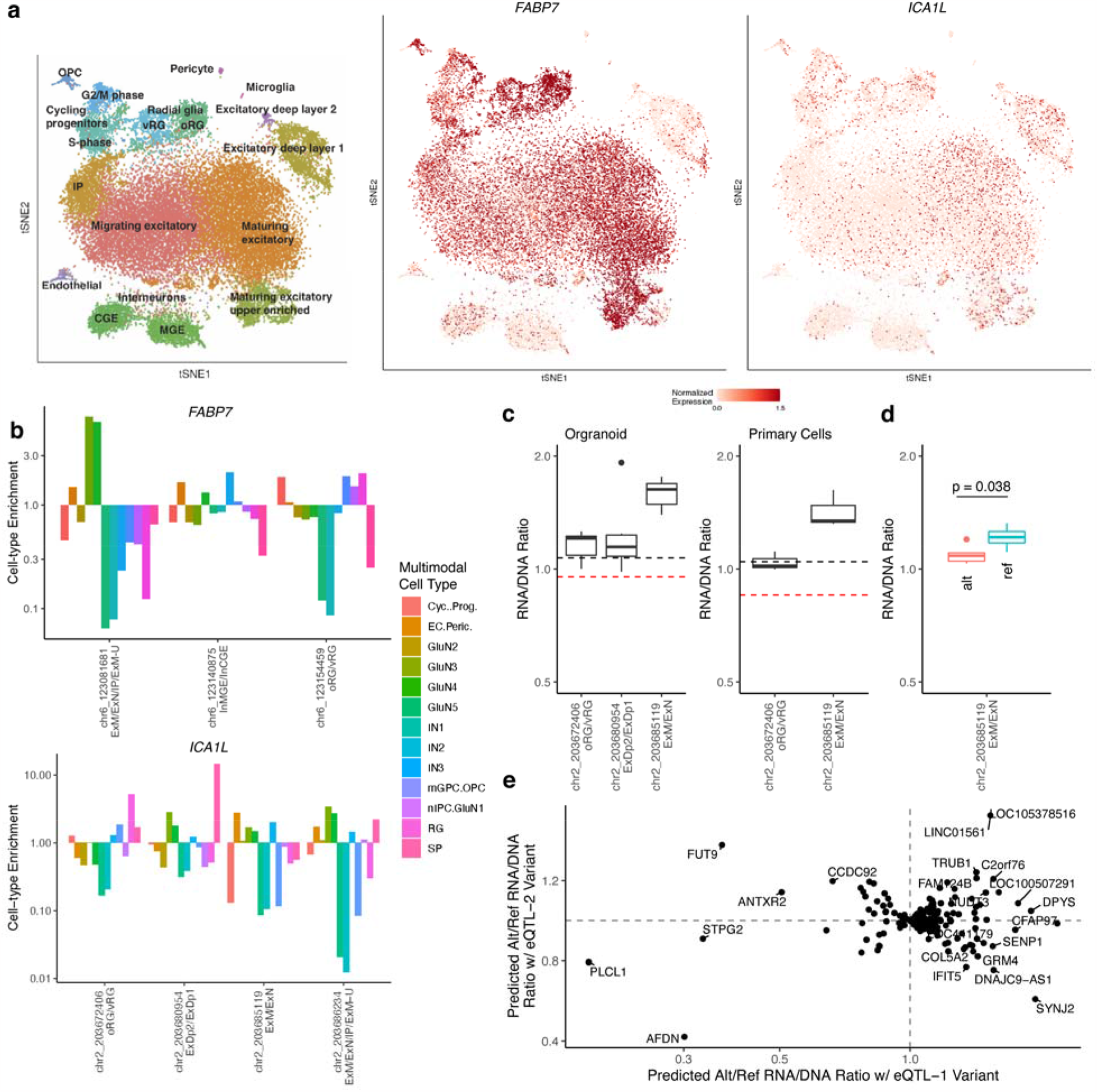
*FABP7* and *ICA1L* exhibit cell-type specific regulation. **a**. *FABP7* (middle) and *ICA1L* (right) are expressed in many cell types in the developing brain (cell type labels shown on left). **b**. For each eQTL associated with *FABP7* (top) and *ICA1L* (bottom) cells in which the respective eQTL variant is accessible in are enriched for the matching cell type in the labeled scRNA-seq portion of jointly profiled multiome scRNA/ATAC data. **c**. MPRA data from brain organoid (left) and primary cells (right) that overlapped overlapped *ICA1L* eQTLs all showed RNA/DNA ratio at or above the median of positive controls (black dashed line) and above the median for negative controls (red dashed line) indicating that these variants occur in functional regulatory regions. **d**. For one *ICA1L* eQTL variant, both the reference (red) and alternative (blue) alleles were tested in an MPRA finding a significant difference in RNA/DNA ratio. **e**. A machine learning model predicts that the ratio between alternative and reference alleles of RNA/DNA ratio for genes with cell-type divergent eQTLs is often independently changed by both eQTLs.

A similar enrichment for corresponding cell types was observed for *ICA1L* eQTLs, four of which could be tested using multiome data (Figure 4b, bottom). For three of these eQTLs the variant site in question had previously been tested in an MPRA experiment conducted in cortical organoids and primary cortical cells.^17^ In all three, the MPRA activity (RNA/DNA ratio) was greater than or equal to positive controls, and always higher than negative controls (Figure 4c), indicating that these are functional variants. Furthermore, for one variant both the reference and alternative allele were tested in the MPRA, and there was a significant difference in the RNA/DNA ratio between alleles (Figure 4d). This data validates our prediction of cell-type specific regulatory variants for *ICA1L*.

Due to the low overlap between regions tested in MPRAs and eQTLs, we expanded our analysis using a machine learning model trained on MPRA data.^17^ We predicted RNA/DNA ratios for both the reference allele and alternative allele for the variant site for each eQTL and found that among genes with cell-type divergent eQTLs, 23% had an eQTL that was predicted to be functional (defined as an absolute log-fold change in the ratio between alleles greater than 0.2, the threshold for significance at a false discovery rate < 0.1 in the original MPRA study). Of these, 16% had a second eQTL that was annotated to a different hierarchical cell type and was also predicted to be functional (Figure 4e). Thus, machine learning provides further support for the presence of many genes with cell-type divergent eQTLs regulating different components of their expression across the developing brain.

## Discussion

Hierarchical cell type annotation enabled the identification of eQTLs with cell-type divergent regulation of genes. Crucially, it showed clear advantages over using a non-hierarchical approach or simply overlapping eQTLs with annotated enhancers or regulatory regions. Additionally, our approach gives a continuous score to each eQTL across every level of the cell type tree rather than just a binary annotation for each cell type. By not selecting a particular resolution of cell type and not relying on any prior clustering of cells, our approach allows for a more flexible view of annotation. It also leverages relationships between cell types to annotate eQTLs.

Our observation that changes in accessibility around eQTL variants link to cell-type specific expression of genes, potentially driven by altered transcription factor binding motifs, agrees with the currently held views of gene pleiotropy. However, lingering questions on the combinatorial operation of enhancers and pleiotropic enhancers were not addressed in this study. With a larger and higher read depth set of single-cell sequencing data it would be feasible to use our same approach to study multiple eQTL variants at the same time. Unfortunately, current scATAC-seq coverage is too low.

The hierarchical model we propose can be readily extended to more complex representations of cell types beyond cell type trees. While we directly encode cell type trees as parent nodes each with two children nodes, our framework allows for the cell types to be any graph. This includes more than two descendant cell types for each parent, as well as cell types that are ambiguously descended from multiple parents. Furthermore, edges can be weighted to represent the probability of two cell types being related or descended from each other. Overall, a graph provides a flexible model of cell type relationships.

Finally, while this study focused on eQTLs with the goal of examining gene pleiotropy, hierarchical annotation can be applied more generally. Our model could be used to hierarchically annotate any genomic regions, including bulk-derived regulatory elements, GWAS hits or other noncoding variants, and more. While each of these different applications would require a careful study of the corresponding data to build the correct network representation, the general framework we have proposed is universal.

## Methods

### Hierarchical model construction

The hierarchical model was implemented as an extension of the CellWalkR (version 0.99.1).^18^ First, a cell type hierarchy, which is typically represented by a tree in which the leaf nodes are cell types, is converted into a symmetrical (2*n*-1)-by-(2*n*-1) adjacency matrix where *n* is the number of cell types in the tree, representing the total number of leaf and internal nodes in a tree. Each parent-child relationship in the tree is given a value of one in the adjacency matrix, and all other values are set to zero. Next, a (2*n*-1)-by-*c* matrix, where *c* is the number of cells, is constructed by padding the *n*-by-*c* label-cell matrix constructed by CellWalkR from cell type labels and scATAC-seq data with an (*n*-1)-by-*c* matrix of zeros, representing that no internal tree nodes are directly connected to cells. Finally, a symmetrical (2*n*-1+*c*)-by-(2*n*-1+*c*) matrix is constructed by appending the (2n-1)-by-(2n-1) matrix, the (2*n*-1)-by-*c* matrix, and transposed *c*-by-(2*n*-1) copy of that matrix, and the *c*-by-*c* cell-cell matrix generated by CellWalkR to each other.

Once the matrix is constructed, it is used in place of the standard matrix generated by CellWalkR for all downstream functions. A random walk with random restarts as implemented in CellWalkR’s *randomWalk* function calculates how much information flows from each node to each other node. In particular, this includes the probability that a walk starting at each cell node ends at each cell type node as well as each internal node representing portions of the cell-type hierarchy. Bulk data is annotated to each cell type node and each internal node using CellWalkR’s *labelBulk* function.

### Cell-type eQTL scoring

To score eQTLs from the developing brain we first built a hierarchical model for cell types in the developing brain. The cell type hierarchy from Polioudakis et al.^15^ was encoded as described above with 16 leaf nodes (corresponding to cell types) and 15 internal nodes. We then ran our extended version of CellWalkR using cell type marker genes from Polioudakis et al. and scATAC-seq from Ziffra et. al^19^ with the “logFC” option for marker genes corresponding to cell types and the *mapSnapATACToGenes* function with the “whichMat” option set to “gmat” to generate label-cell edges and *computeCellSim* to generate cell-cell edges. We tuned edge weights using the *tuneEdgeWeights* function with steps set to three and found that 100 was the optimal setting of the edge weight parameter. We ran the *walkCells* function with the optimal edge weight to calculate a final influence matrix.

161,929 fine-mapped eQTLs were taken from Wen et al.^9^ and lifted from hg19 to hg38 using liftOver.^21^ They were then mapped to each of the 16 cell types and 15 internal nodes using the *labelBulk* function. This resulted in a vector of 31 scores for 11,765 eQTLs across 4,137 genes, corresponding to the set of eQTLs that could be scored based on the scATAC-seq data. Of these, 5,889 mapped to at least one node with a label score greater than two, indicating high specificity. For downstream analyses, for each eQTL we then looked to see if it was significant in more than one hierarchical cell type. As a conservative annotation approach, in cases where an eQTL was significant in cell types that were ancestors of each other in the hierarchy, we only considered the highest level (i.e. least specific) annotation, and removed any more specific annotations.

For comparison to a non-hierarchical annotation we ran a standard version of CellWalkR with the same data and options, and found that 100 was the optimal setting of the edge weight parameter. This model scored the same 11,765 eQTL SNPs across 4,137 genes this time with a vector of 16 scores, each for each cell type. We used Used a label score threshold of two for high specificity and one for low specificity. To determine if each eQTL variant overlapped with a cell type specific peak or enhancer, we downloaded annotated regions from Ziffra et al.^19^

We embedded the length 31 label score vectors for each eQTL into two dimensional space using the UMAP method (as implemented in the uwot package version 0.1.10 in R) with default parameters.^22^ For each gene, we considered it to have divergent eQTLs if at least two eQTLs mapped to that gene which were not ancestors of each other in the original hierarchy, had a euclidean distance of at least 40 between their length 31 cell type vectors (to ensure they are not highly correlated) and had a euclidean distance of at least 8 in UMAP space (to ensure the eQTLs don’t correspond to similar cell types).

### Functional validation

scRNA-ATAC multiome data was downloaded from Trevino et al.^16^ We calculated the entropy of each gene from the scRNA-seq portion of the data as described in Kannan et al.^23^ We assigned multiome cells to each eQTL if that eQTL’s variant site was accessible in that cell but no other eQTL variant for the same gene was accessible. We then calculated the log-fold change in mean expression and a p-value (using a two tailed Wilcoxon test) for the gene for each eQTL between cells assigned to that eQTL and those not assigned to any eQTL. A false discovery rate was calculated for the *p*-values using the Benjimani-Hochberg procedure. We filtered for highly expressed genes as those that had at least 100 reads. Cell type enrichment was calculated by taking the scRNA-based cell type assigned to each cell and for each eQTL and computing the fraction of cells assigned to that eQTL that are of that cell type over the fraction of cells not assigned to that eQTL that are that cell type.

Transcription factor (TF) binding motif disruption for each eQTL variant was calculated using motifbreakR (version 2.14.2) with default parameters.^24^ A TF was considered expressed in a cell type if it had at least 100 reads per million mapped reads in that cell type in the Polioudakis et al scRNA-seq data.^15^ The TF by gene heatmap and TF by TF heatmap were generated using the *heatmap*.*2* function in R with the “symm” variable set to TRUE. The first is trimmed to only show counts greater than five, and the second to only show counts greater than two. Expression UMAPs for *FABP7* and *ICA1L* were generated from CoDEx Viewer.^15^

To validate the *FABP7* locus, known enhancer data was downloaded from FANTOM5^25^ and candidate cell type specific regulatory elements were downloaded from Deng et al.^17^ Massively Parallel Reporter Assay (MPRA) data for cerebral organoid and primary fetal cortical cells were also downloaded from Deng et al. The machine learning model from that paper was run on each eQTL variant to predict an RNA/DNA ratio for each reference and alternative allele.

## Availability of Data and Materials

No datasets were generated during the current study. All analyzed data is publicly available or available by request from the corresponding publications.

