## Supplemental Figures for "Hierarchical annotation of eQTLs enables identification of genes with cell-type divergent regulation"

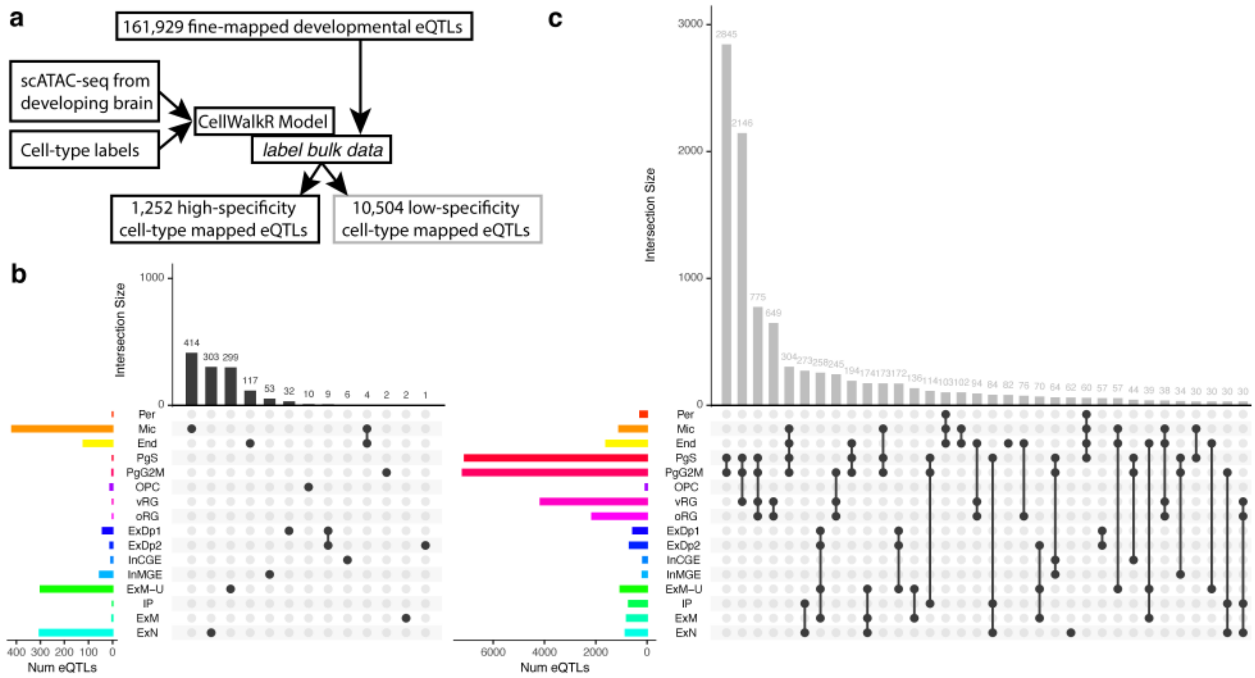


Supplementary Figure 1. **Mapping non-hierarchical cell types to eQTLs a.** A CellWalkR model was built using scATAC-seq data and cell-type labels from the developing brain to label fine-mapped developmental brain eQTLs to cell types. **b.** High-specificity cell-type mapped eQTLs generally map to a single cell type, but they are biased towards very distinct or common cell types. **c.** Low-specificity cell-type mapped eQTLs map to a broader set of cell types, but often to multiple closely related ones.

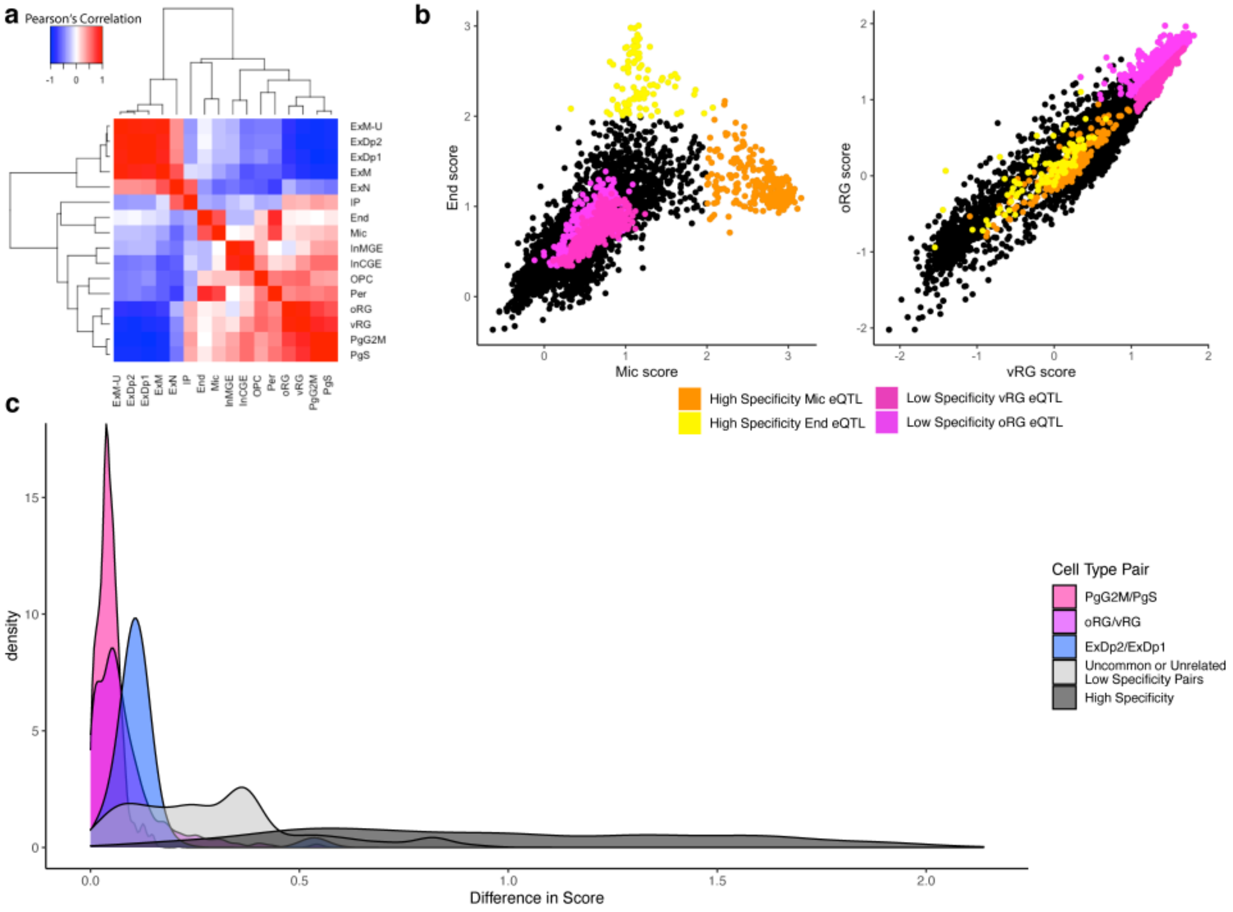


Supplementary Figure 2. **Label scores for cell types in the developing brain can be highly correlated. a.** A clustered heatmap of correlations of label scores across cells shows that some labels are highly correlated **b.** Label scores between distinct cell types like Microglia (Mic) and Endothelial (End) cells are highly correlated overall, but show high specificity for those cell types (left panel). Label scores for similar cell types like outer Radial Glia (oRG) and ventricular Radial Glia (vRG) are highly correlated and also show low specificity for those cell types (right panel) **c.** The distributions of the differences between the top scores for the three most common cell type pairs for highly related cell types in low-specificity cell-type mapping as well as for unrelated pairs and for highly specific labels indicate that label scores for highly related cell types are often very similar compared to unrelated cell types.

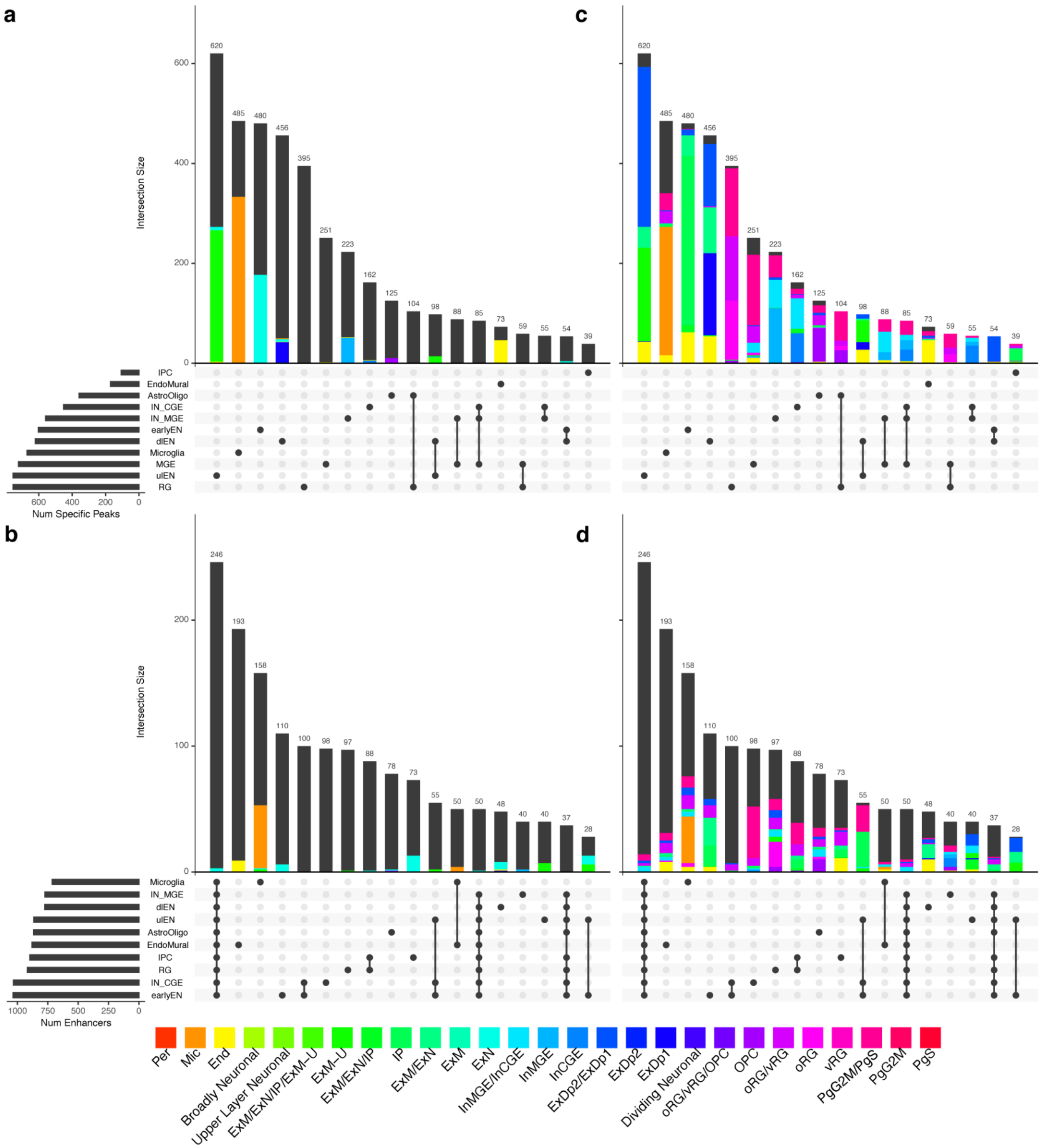


Supplementary Figure 3. **eQTL mapped cell-types overlap with cell-type specific peaks and enhancers.** The overlap of eQTLs with **a.** cell-type specific peaks **b.** and predicted enhancers colored by mapped non-hierarchical cell types and the same annotations colored by mapped hierarchical cell types (**c** and **d**). We observe a greater than four-fold increase overall in the number of eQTLs in cell-type specific peaks that overlap hierarchical labels vs non-hierarchical labels (4,091 vs 1,009), with 93.2% of eQTLs in cell-type specific peaks overlapping a hierarchical label. We see a similar increase in cell-type specific enhancer coverage (those assigned to fewer than 3 cell types, 531 vs 131, out of 1,417 such enhancers), with only a minimal increase in cell-type mapping to broad enhancers (those assigned to all cell types, a gain of 9 out of 246 such enhancers).

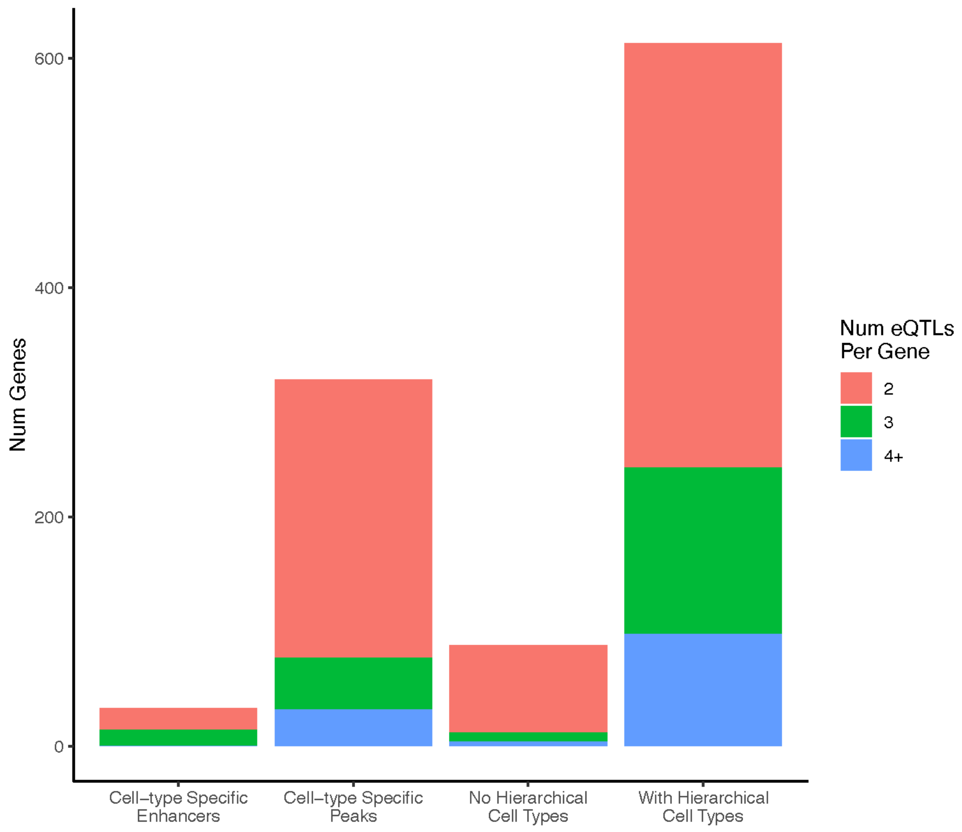


Supplementary Figure 4. **Hierarchical annotation of cell types increases the number of detectable genes with cell-type divergent eQTLs.** Using alternative methods for annotating eQTLs such as overlap with cell-type specific enhancers, overlap with cell-type specific peaks, or annotation with non-hierarchical cell types detects mich fewer genes with cell-type divergent eQTLs than annotation with hierarchical cell types.

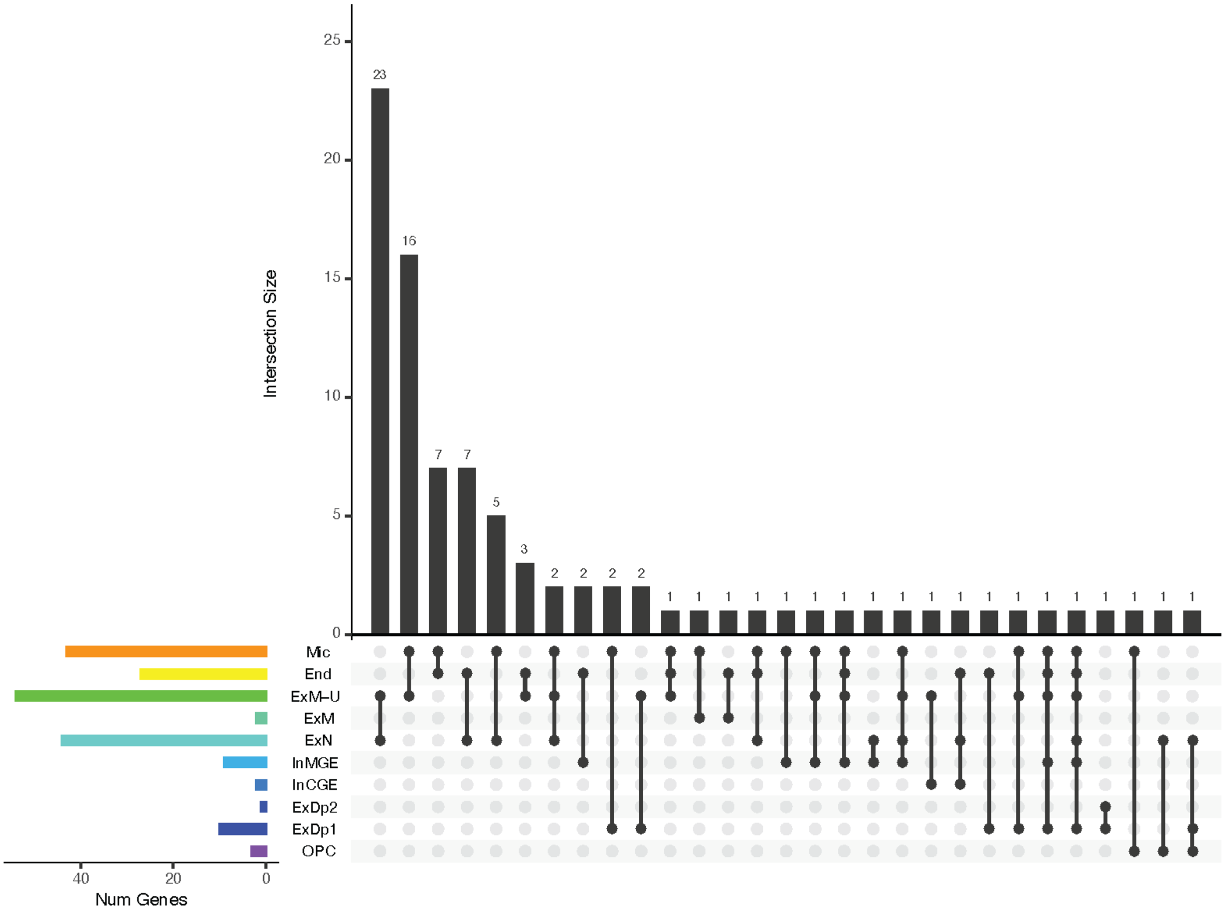


Supplementary Figure 5. **Divergent non-hierarchical cell types for eQTLs across genes.** Among genes with cell-type divergent eQTLs detected in non-hierarchical cell types, the majority of divergent annotations included one of the two most distinct cell types (Microglia and Endothelial cells), with few combinations covering other types of cells.

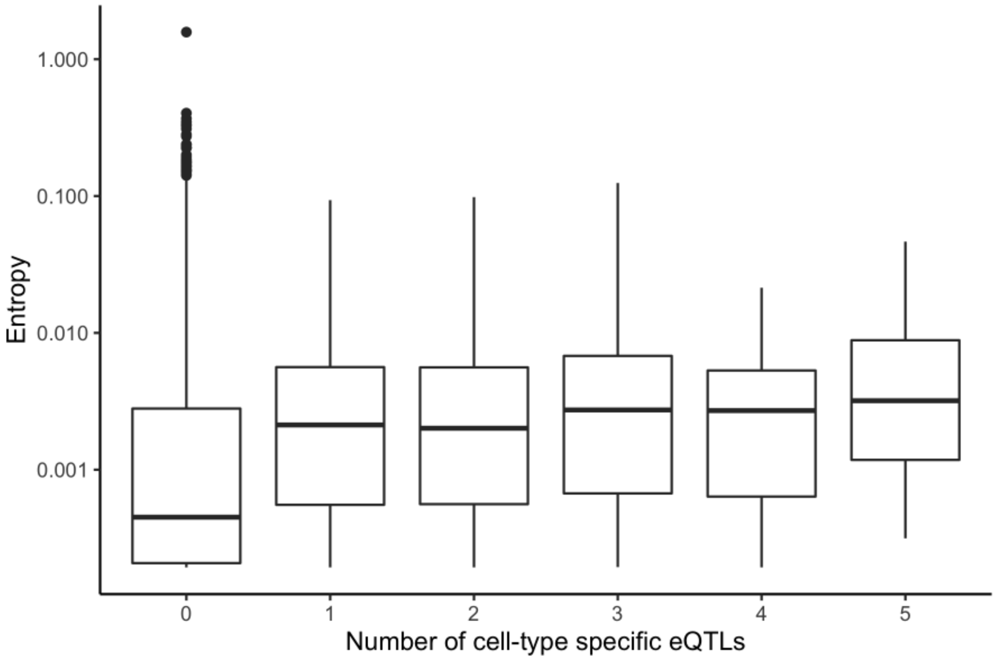


Supplementary Figure 6. **Entropy increases with the number of cell-type specific eQTLs.** Genes exhibit more entropy in their expression across cell types when they are annotated with more cell-type specific eQTLs, indicating that more eQTLs allow for more cell-type specific expression.

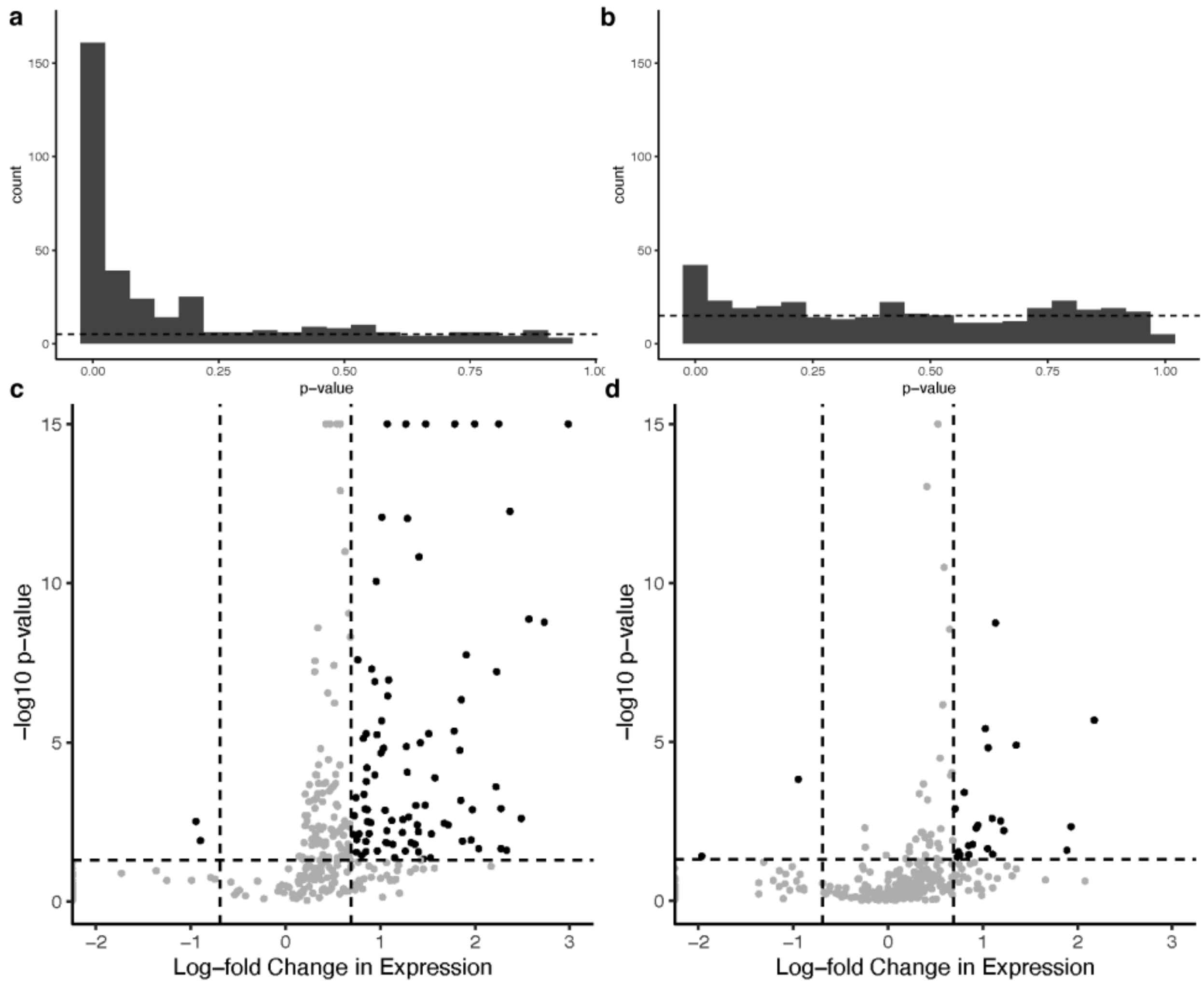


Supplementary Figure 7. **Genes with cell-type divergent eQTLs are differentially expressed. a.** Distribution of *p*-values for differential expression between cells that have only the most frequently accessible eQTL variant for that gene accessible versus those that do not. **b.** Distribution of *p*-values for differential expression between cells that have only the second most frequently accessible eQTL variant for that gene accessible versus those that do not. In both cases there is an enrichment for significant *p*-values. Panels **c.** and **d.** show the same genes as plotted as volcano plots, identifying the 30 genes that have significant log-fold changes in expression across multiple eQTLs.

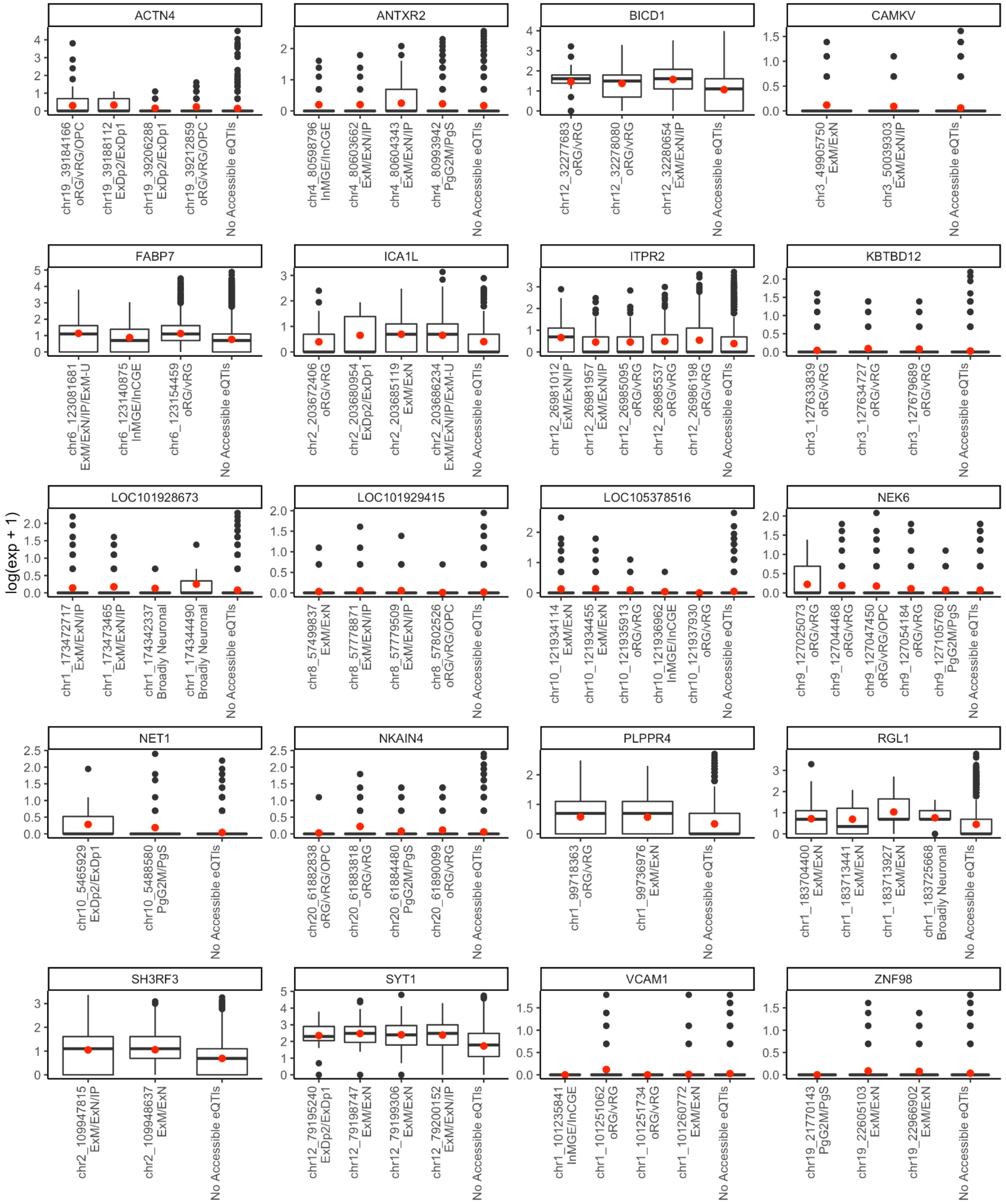


Supplementary Figure 8. **Multiome expression data for eQTLs.** For highly expressed genes, boxplots of the RNA-seq read counts of cells in which the given eQTL is accessible in ATAC-seq in multiome scRNA/ATAC-seq data, with mean counts shown with red dot.

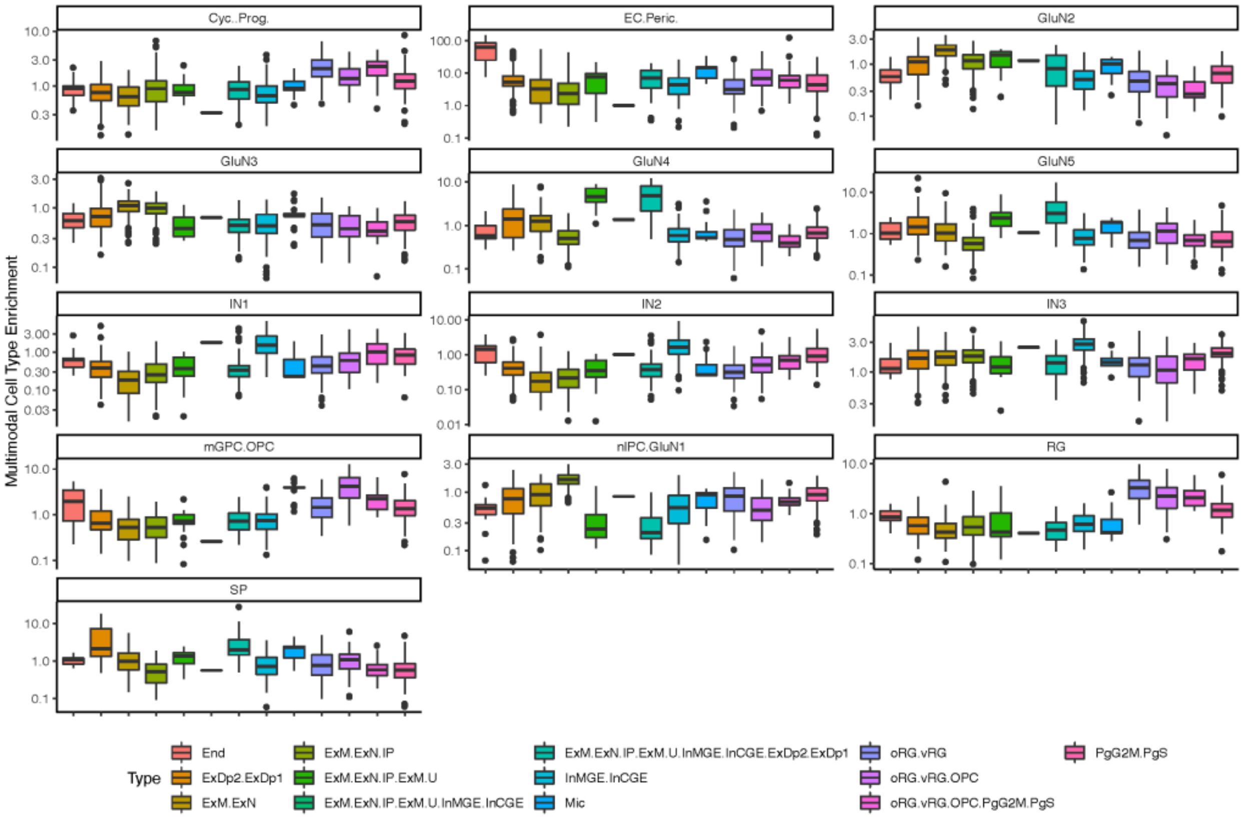


Supplementary Figure 9. **Multiome cell-type enrichment for hierarchical cell types.** Across cell-types annotated in the RNA-seq portion of multiome scRNA/ATAC-seq data (shown in panel titles), hierarchical cell-type annotated eQTLs that are accessible in the corresponding scATAC-seq data for the same cell are enriched for similar cell types.

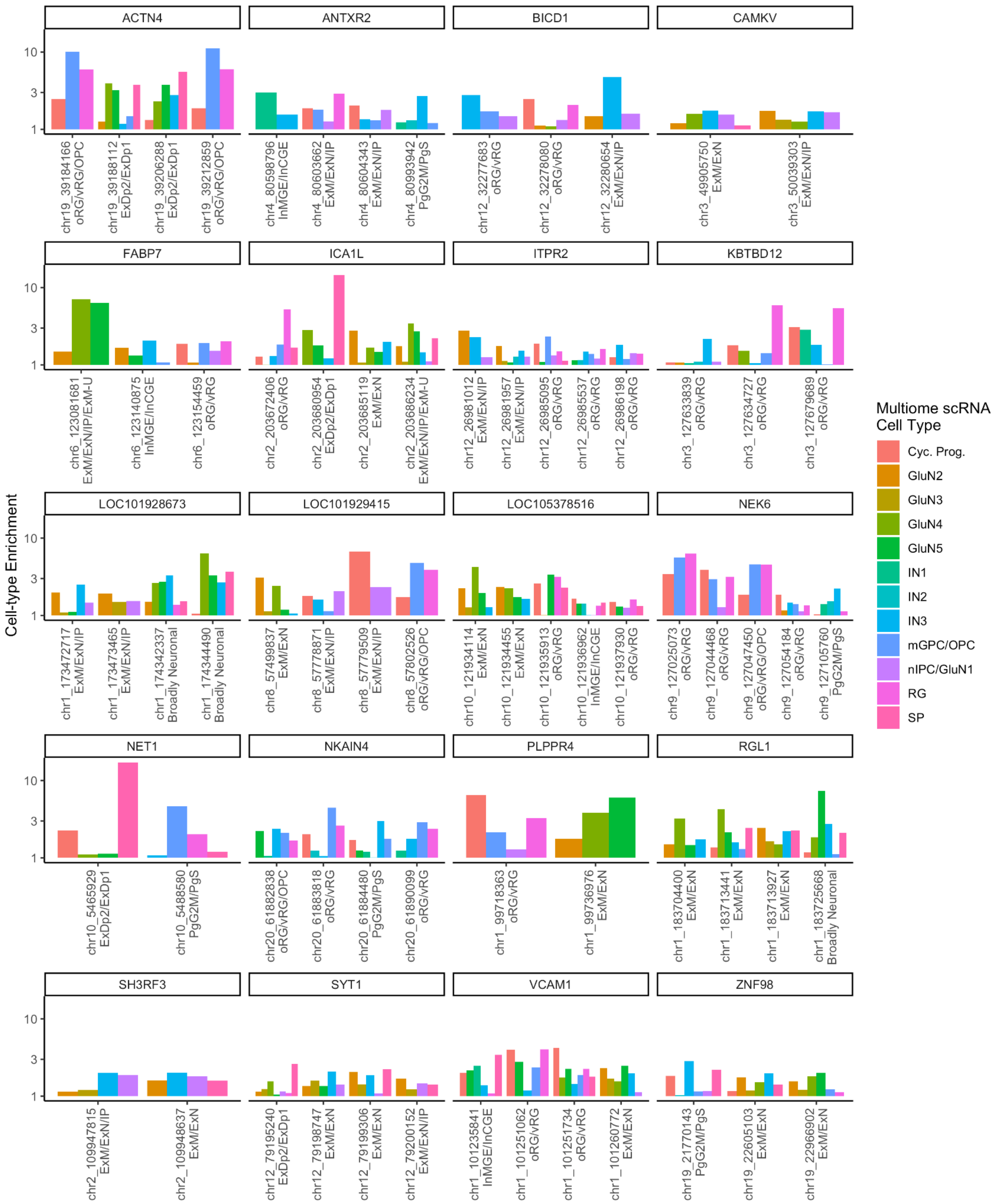


Supplementary Figure 10. **Multiome cell-type enrichment for eQTLs.** For highly expressed genes, enrichment of the cell type annotated in scRNA-seq of cells in which the given eQTL variant is accessible in ATAC-seq in multiome scRNA/ATAC-seq data (only positive enrichment shown).

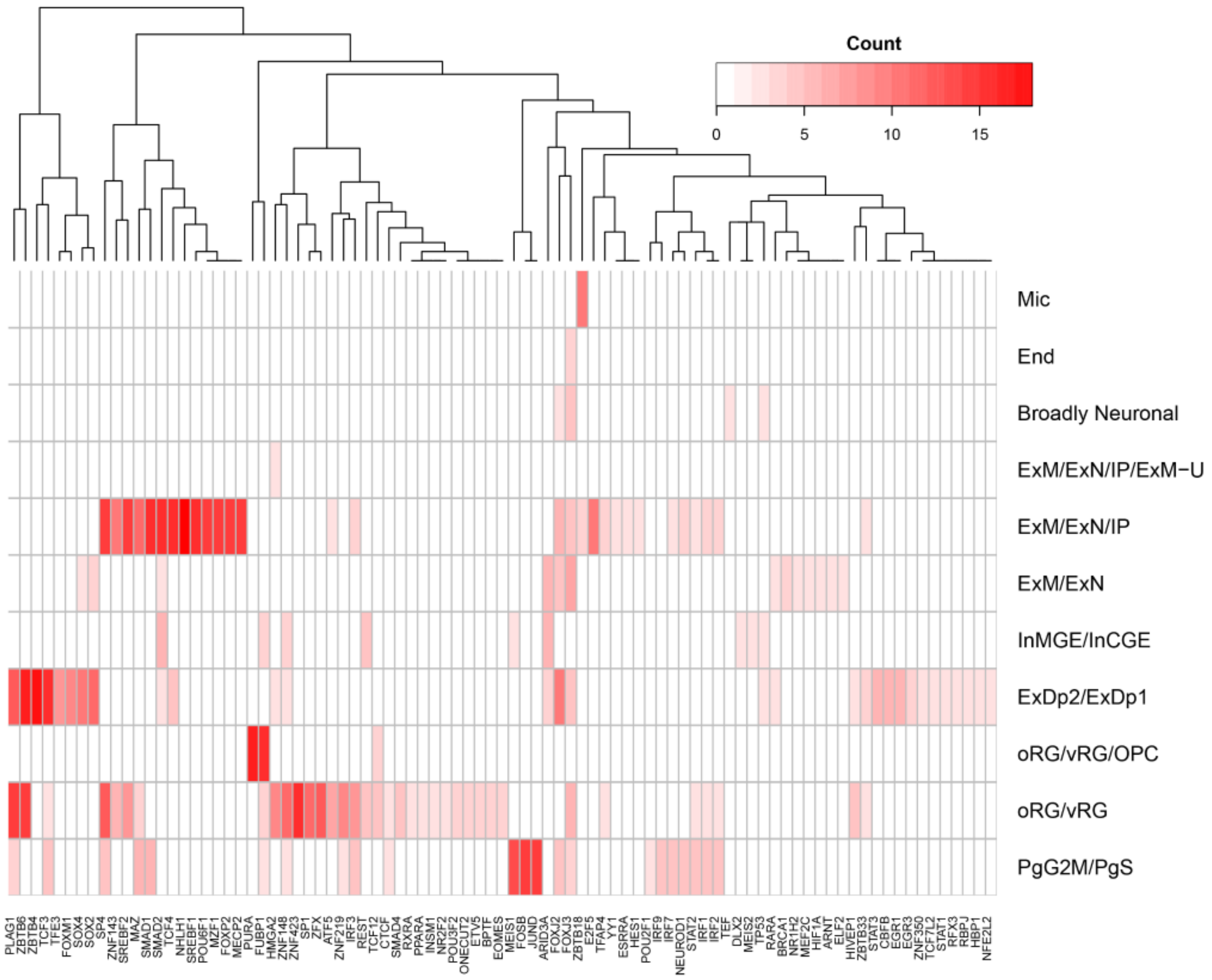


Supplementary Figure 11. **Cell-type specific TF Binding.** A heatmap showing how frequently transcription factor (TF) binding sites were predicted to be significantly disrupted by MotifBreakR by eQTLs associated with each hierarchical cell type.

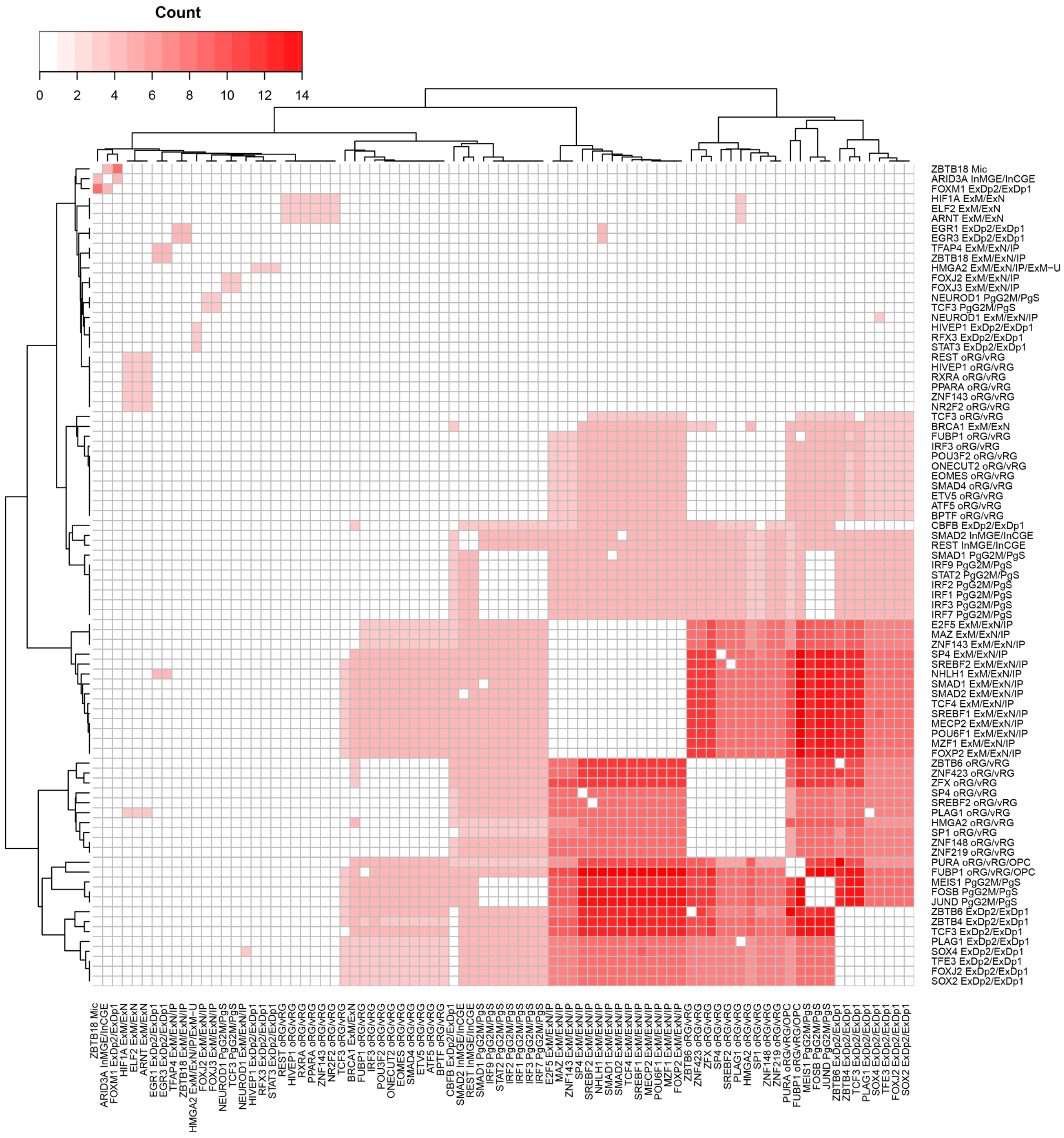


Supplementary Figure 12. **Co-occurring TFs.** A heatmap showing how frequently pairs of transcription factor (TF) binding sites were predicted to be significantly disrupted by MotifBreakR by eQTLs associated with different hierarchical cell types for the same gene.

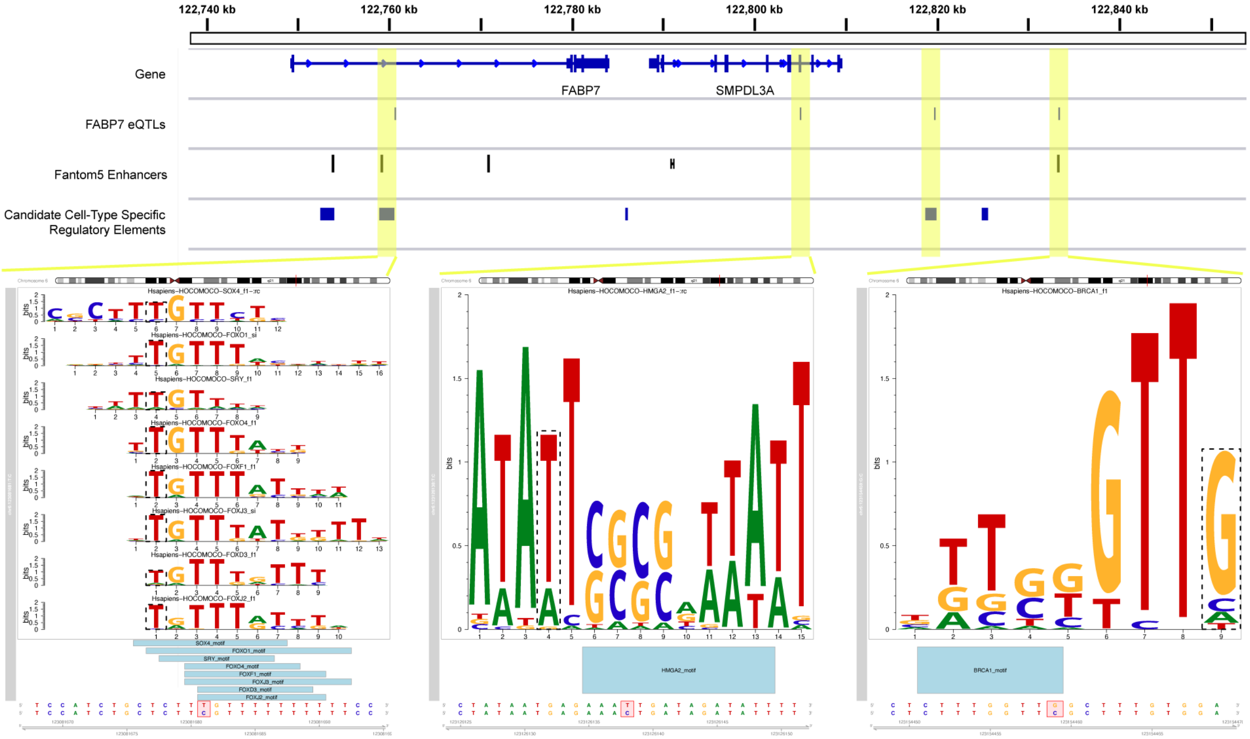


Supplementary Figure 13. **The *FABP7* Locus.** Of the four eQTLs (highlighted in yellow) that are linked to *FABP7*, two lie in known enhancers, two lie in predicted cell-type specific regulatory elements, and three disrupt the binding sites of different transcription factors.
